# The cys-loop ligand-gated ion channel gene superfamily of the Colorado potato beetle, *Leptinotarsa decemlineata*

**DOI:** 10.1101/2025.01.13.632561

**Authors:** Dongxu Chen, David J. Hawthorne

## Abstract

**Background:** The insect cys-loop ligand-gated ion channel (cysLGIC) superfamily includes nicotinic acetylcholine receptors (nAChRs), γ-aminobutyric acid (GABA) receptors, glutamate- or histamine-gated chloride channels (GluCls and HisCls), pH-sensitive chloride channels (pHCls) and several other functionally uncharacterized receptors. Several of these receptors are target sites of neonicotinoids and other insecticides. Characterizing sequences of cysLGIC genes can facilitate the study of functional expression of subunits allowing insecticide/receptor interaction research, and can promote molecular resistance monitoring tools development. The Colorado potato beetle (CPB) is an agricultural pest that threatens the production of solanaceous crops. Although this insect shows frequent evolution of insecticide resistance, its cysLGIC superfamily is not well characterized.

**Results:** Twenty-two candidate CPB cysLGIC subunit genes were identified, and the functional regions of their protein sequences were annotated. CPB possesses 22 candidate cysLGIC subunit genes such as nAChR α4, nAChR α6, RDL, and GluCl subunits, with similar sequence, structure, and alternative exon use as that in other insects. RNA A-to-I editing was observed of nAChR α6. Two copies of the pHCl subunit gene were identified, the first duplication of this gene observed in insects.

**Conclusion:** The number of cysLGIC superfamily genes is similar to that of other insect species. Alternative splicing and RNA editing conserved in insect species were also identified in expected subunits, potentially contributing to structural and functional diversity of the receptor. Evidence of naturally truncated nAChR α4 and duplicated pHCl was observed, which invites future validation.

## Background

Members of the cys-loop ligand-gated ion channel (cysLGIC) superfamily mediate both fast excitatory and inhibitory synaptic transmission in the nervous system of vertebrates and invertebrates. The cysLGIC are formed by the assembly of five subunits that share similar structural features: a large extracellular N-terminal region containing the signature cys-loop (two cysteines separated by 13 amino acids and bonded by disulfide bonds), ligand-binding motifs (loops A–F), and four transmembrane (TM) domains that form the ion-selective pore [1] (Figure 1). Insect cysLGICs include the cation-permeable nicotinic acetylcholine receptors (nAChRs) [2], the γ-aminobutyric acid (GABA) receptor with both cationic and anionic (chloride) channel properties [3], the glutamate- or histamine-gated chloride channel (GluCl and HisCl) [4, 5], the pH-sensitive chloride channel (pHCl) [6], and the insect group1 subunits and several other functionally uncharacterized receptors [7] (Table 1, Table S1). In addition to their roles in the insect nervous systems, these receptors are of particular interest as target sites of widely used insecticides. For example, nAChRs are target sites of the neonicotinoid and spinosyn insecticides, and GABA receptors are target sites of cyclodienes and the phenyl pyrazoles. According to the insecticide resistance action committee (IRAC) mode of action classification, members of the cysLGIC superfamily are target sites of seven insecticide groups (Table 1).

**Figure 1.**
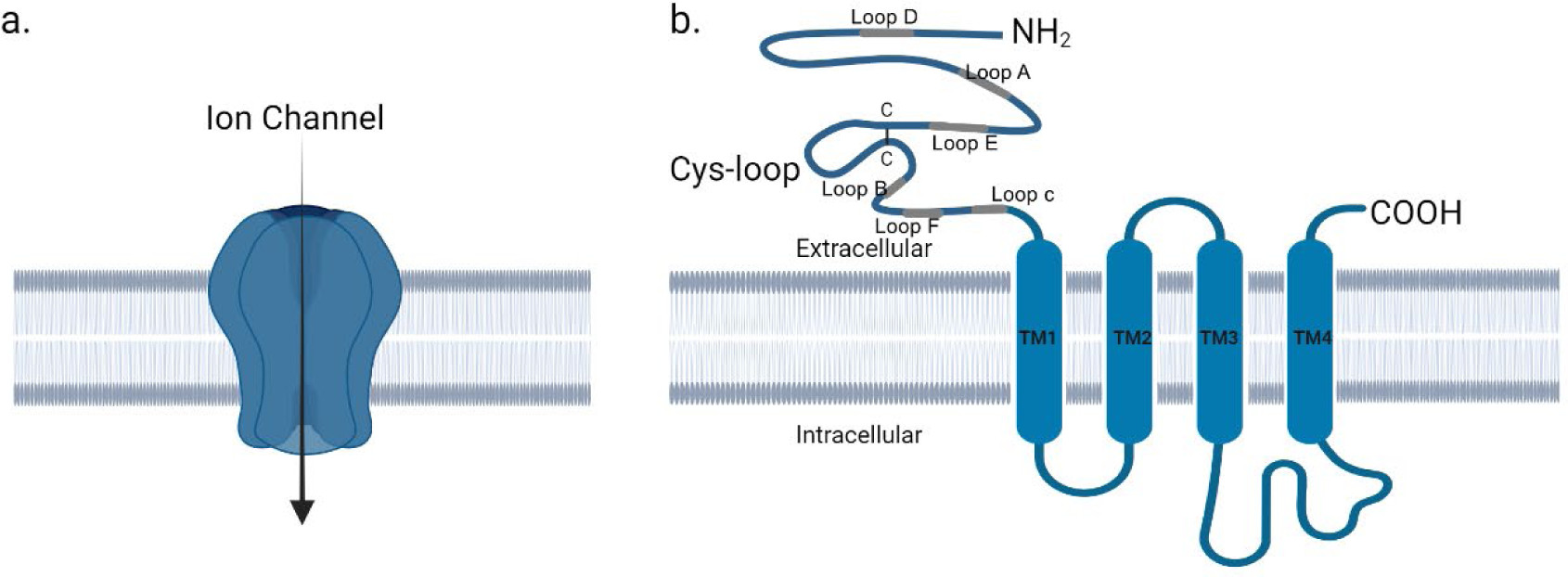
Structure of a typical cys-loop receptor. a). The pentameric receptor with ion permeability; b). Topology of a cys-loop receptor subunit. The extracellular domain contains the signature cys-loops and Loops A-F (Gray shaded region) for ligand binding. Sequences connecting transmembrane domains 1 to 3 are usually short and can be longer between TM3 and TM4.

**Table 1.** Sub-category of insect cys-loop ligand-gated ion channels and their role as insecticide target sites.

| Receptor Name/Description | Endogenous ligand | Nomenclature of subunits | Targeted Insecticide Group |
| --- | --- | --- | --- |
| nicotinic acetylcholine receptor | Acetylcholine (ACh) | $\alpha$ (possessing two consecutive cysteines in loop C) and $\beta$ | Group 4: neonicotinoid, Sulfoximines, etc.<br>Group 5: Spinosyns<br>Group 14: Nereistoxin analogues<br>Group 32: GS-omega/kappa HXTX-Hv1a peptide |
| γ-amino butyric acid-gated anion channels | γ-amino butyric acid (GABA) | RDL, LCCH3 and GRD | Group 2: Cyclodiene<br>Organochlorines & Fiproles<br>Group 30: Meta-diamides & Isoxazolines |
| Glutamate-gated chloride channel | Glutamate | GluCl | Group 6: Avermectins & Milbemycins) |
| Histamine-gated chloride channel | Histamine | HisCl |  |
| pH-sensitive chloride channel | unclear | pHCl |  |
| Insect group 1 cysLGIC. Initially annotated in *D. melanogaster* as CG6927, CG7589 and CG11340 | unclear | CLGC |  |
| Ungrouped subunits. Initially annotated in *D. melanogaster* as CG8916 and CG12344 | unclear | 8916 or 12344 |  |

Subunits are labelled with a one-letter genus code plus a three-letter species code and the subunit types. For example, the nAChR
$\alpha 6$ subunit in *Tribolium castaneum* was denoted as Tcas\_ $\alpha 6$ and the RDL ortholog of CPB is Ldec\_RDL.

Insect nAChRs respond to acetylcholine (ACh) and mediate fast excitatory transmission at cholinergic synapses in the central nervous system (CNS). They involve processing sensory signals, locomotion, and cognitive processes [8]. Insect nAChR subunits are of two types, α (characterized by the two consecutive cysteines in Loop C) and β [2]. Most insect nervous nAChRs are hetero-pentameric, containing both α and β subunits, while in some rare cases, particular α subunits can form homomers [9]. Imidacloprid acts at the ACh binding site (the orthosteric site) on insect nAChRs, thereby activating the integral cation-permeable ion channel of the receptor [10] (Figure 1). GABA receptors are widespread in the insect CNS, mediating inhibitory and excitatory neurotransmission. The first insect GABA subunit gene was identified from a *D. melanogaster* strain *r*esistant to *d*ie*l*drin (a cyclodiene insecticide), named *RDL* [11]. GluCls have a wide range of functions in invertebrates’ nervous systems associated with locomotion, feeding, and sensory inputs [4]. The first insect GluCl subunit cDNA was cloned from *D. melanogaster* [12]. Most insect species, including *A. mellifera, T. castaneum,* and *N. vitripennis*, have a single GluCl gene. Histamine is the primary neurotransmitter of arthropod photoreceptors, and these histamine-gated chloride channels likely play a role in vision, although their function remains unclear [12, 13]. The pH-sensitive chloride channel was initially identified in *D. melanogaster*. While its native ligand has not yet been identified, its chloride channel is opened in response to shifts in extracellular pH [6]. Insects possess 1 - 3 group1 subunits. In early studies, these subunits were labeled by numbers, consistent with the first discovered *D. melanogaster* orthologs (CG6927, CG7589, and CG11340); later, subunits belonging to this group, were labeled as CLGC, standing for Cys-loop Ligand Gated ion Channel [14]. Finally, there are two functionally uncharacterized subunits, initially annotated as CG8916 and CG12344, in the *D. melanogaster* genome assembly.

Insect species typically possess 21-26 cysLGIC subunit genes [7, 14–20], except for two cockroach species showing over 30 members, mainly because of expansion of nAChR β subunits [21]. These numbers are relatively small compared with other arthropod and nematode cysLGIC superfamilies; for example, the spider *Pardosa pseudoannulata*, the spider mite *Tetranychus urticae*, and the nematode *Caenorhabditis elegans* have 34, 29, and 102 cysLGIC genes respectively [22–24].

Genomic analysis of these proteins in pest insects has several benefits. Variation in cysLGIC receptor genes conferring target site insensitivity have been identified in insects (Table 2). Characterizing this variation provides the basis for molecular resistance monitoring. Similarly, sequence knowledge allows the development of nucleic acid-based control methods such as RNA interference.

**Table 2.** Target site insensitivity mutations on cys-loop receptors.

| Subunits | Mutations | Species | Resistance to | Reference |
| --- | --- | --- | --- | --- |
| nAChR β1 | R81T | *Myzus persicae* | Neonicotinoid | [25, 26] |
|  |  | *Aphis gossypii* | Neonicotinoid | [27, 28] |
|  | V101I | *Myzus persicae* | Neonicotinoid | [25] |
|  | L80S | *Aphis gossypii* | Imidacloprid | [29] |
|  | A58T&R79E | *Bemisia tabaci* | neonicotinoid | [30] |
| nAChR α6 | G275E | *Frankliniella occidentalis* | Spinosad | [31] |
| RDL | A301S/G | *A. aegypti* | Fiproles | [32] |
|  |  | *Nilaparvata lugens* | Fipronil | [33, 34] |
|  |  | *Blattella germanica* | Fipronil and dieldrin | [35, 36] |
|  | T350M<br>T350M+A301 G | *D. simulans* | fipronil | [37] |
| GluCl | L315F | *M. domestica* | ivermectin | [38] |
|  | L319F | *B. mori* | ivermectin | [39] |

Characterizing cysLGIC subunit sequences also aides the study of subunit function, including the expression of subunits in well-established expression vehicles such as *Xenopus laevis* oocytes and gene editing in multiple insect species [8, 40–42].

The Colorado potato beetle (CPB) is a damaging pest, threatening the production of solanaceous crops, including potatoes, tomatoes, and eggplants. Intensive insecticide use has led to resistance to cysLGIC targeting insecticides, such as dieldrin [43], organochlorines [44], neonicotinoids [45, 46] and spinosyns [47] in CPB populations. Except for a few cysLGIC genes [48–50], this gene superfamily has not been studied in the CPB despite the importance of insecticides targeting cysLGICs for its control. A recently published chromosome-level CPB genome assembly [51] provides an opportunity to annotate and characterize the cysLGIC superfamily in this species. Here, we report the first systematic study of the cysLGIC gene superfamily in Colorado potato beetles. The protein coding regions of all subunits were successfully identified in the CPB genome. Characterizing the full complement of CPB cys-loop LGIC subunits represents a critical step in identifying key components of the CPB nervous system and pinpointing particular molecular targets underlying responses to insecticides.

## Materials and Methods

### Bioinformatic analysis

To identify potential CPB cysLGIC genes, a tBLASTn [52] search was executed against the chromosome-level CPB genome [51], with all *T. castaneum* and *P. americana* cysLGIC subunit protein sequences as queries. *T. castaneum* was chosen because, among all species with well-characterized cysLGIC, it is most closely related to CPB. *P. americana* was chosen because it has the largest number of insect cysLGIC subunits. Candidate CPB cysLGIC genes were determined based on their identity with queries, particularly in the N-terminal ligand-binding region and the four transmembrane domains. In a second round, *T. castaneum* cysLGIC subunits were mapped to the CPB genome using the protein2genome model in Exonerate [53] to predict the intron and exon boundaries. To find the complete protein coding sequence of each CPB cysLGIC subunit, as well as to validate its expression in CPB, chromosomal hits identified by tBLASTn and Exonerate were extracted and manually assembled to obtain the genomic DNA (gDNA) sequence of each subunit. Then, the assembled gDNA sequences were used as queries to Blastn against a CPB transcriptome (NCBI accession GGNV00000000.1) [54]. Transcript hits with a bit score higher than 900 were collected.

### Sequence Selection for Protein Analysis

Transcripts from each cysLGIC subunit, were translated to their corresponding protein sequences, aligned, and manually screened for completeness of “cysLGIC subunit signature regions” (including the N-terminal ligand-binding motifs and transmembrane domains). Protein sequences proceeding to alignment were selected according to the following rule: for a subunit of which only one of the transcript-translated proteins shows all “signature regions”, the transcript-translated sequence will be selected; if multiple transcript-translated proteins can display all “signature regions”, the longest one was used; For a subunit of which none of the transcript-translated proteins can show complete “signature regions” or no transcript hit has a score higher than 900, a manual search will be executed in the NCBI database to find any of its transcripts, cDNA or protein coding sequence predicted by automated computational analysis.

### Protein sequence alignment and annotation

CPB nAChR subunits and non-nAChR subunits were aligned separately. Protein alignment was conducted by Clustal X2 [55]. The pairwise alignment was set with a gap-opening penalty of 10 and a gap-extension penalty of 0.1, applying the Gonnet 250 protein weight matrix. Sequence alignment was visualized and annotated using JalView version 2.1.1 [56]. Transmembrane domains of protein sequences were predicted by TMHMM - 2.0 (https://services.healthtech.dtu.dk/service.php?TMHMM-2.0). Protein signal cleavage sites were predicted by SignalP 5.0. Genedoc calculated identities/similarities between protein sequences.

### Phylogenetic Analysis

Phylogenetic analyses were run for nAChR and non-nAChR subunits separately. In MEGAX [57], the MUSCLE algorithm aligned cysLGIC subunits of CPB, *D. melanogaster*, *A. mellifera*, *P. americana,* and *T. castaneum* (Additional file 4: Table S4 includes NCBI accession numbers of all sequences, used for phylogenetic analysis, except for CPB), and the phylogenetic tree was then reconstructed using the maximum-likelihood method and Dayhoff matrix-based model with 1000 bootstraps replication. The tree file was visualized and manipulated by TreeViewer [58].

### Alternative splices and post-transcriptional modification identification

Alternative splicing of a subunit gene was identified by comparing its transcript isoforms. In addition, alternative exons can also be evidenced by tBlastn results, as a region of a protein query can hit multiple genomic regions with a reasonably high identity of each hit, and these alternative regions are tandemly arranged in the genome. Pre-mRNA editing was analyzed by comparing gDNA and transcripts of each subunit and their translated protein sequences.

## Results

### Overview of Leptinotarsa cysLGICs

CPB possesses 22 candidate cysLGIC subunit genes, including 11 nAChR subunit genes and 11 non-nAChR subunit genes. Sixteen subunit genes are found on autosomes and six are located on the X chromosome (Table S2). A total of 21 subunit sequences were aligned and subsequently used in phylogenetic analysis (Table S3)[48, 49]

### CPB nAChR subunits

Eleven nAChR subunits were identified in CPB. Based on characteristic features such as the presence or absence of two consecutive cysteines in loop C (Figure 2), ten subunits were categorized as α type and one as β type (Table S1). Most CPB nAChR subunits had amino acid sequence identities over 60% with their *T. castaneum* counterparts (Table 3) except for Ldec_α9 with 43% identity with Tcas_α9. This relatively low identity between nAChR α9 orthologs is not surprising because previous studies found this subunit to be highly variable across species, such as 28% and 20% identity between the *T. castaneum* nAChR α9 and its counterparts in *A. mellifera* and *P. americana*, respectively [14, 21]. In addition, a GEK motif preceding TM2, which is conserved among insect nAChR subunits, was present in all CPB nAChR subunits except Ldec_ α9 (Figure 2). Similarly, replacements of the GEK motif were observed in multiple nAChR subunits of two cockroach species [21], silkworms [59], *N. vitripennis* [16], as well as the nAChR α9 in *T. castaneum*. Ldec_α1 - Ldec_α4 and Ldec_α8 have insertions in Loop F (Figure 2), also seen in other insect species, such as cockroaches [21], parasitoid wasp *N. vitripennis* [16] and the red floor beetle *T. castaneum* [14], which may contribute to interactions with the neonicotinoid, imidacloprid [60].

**Figure 2.**
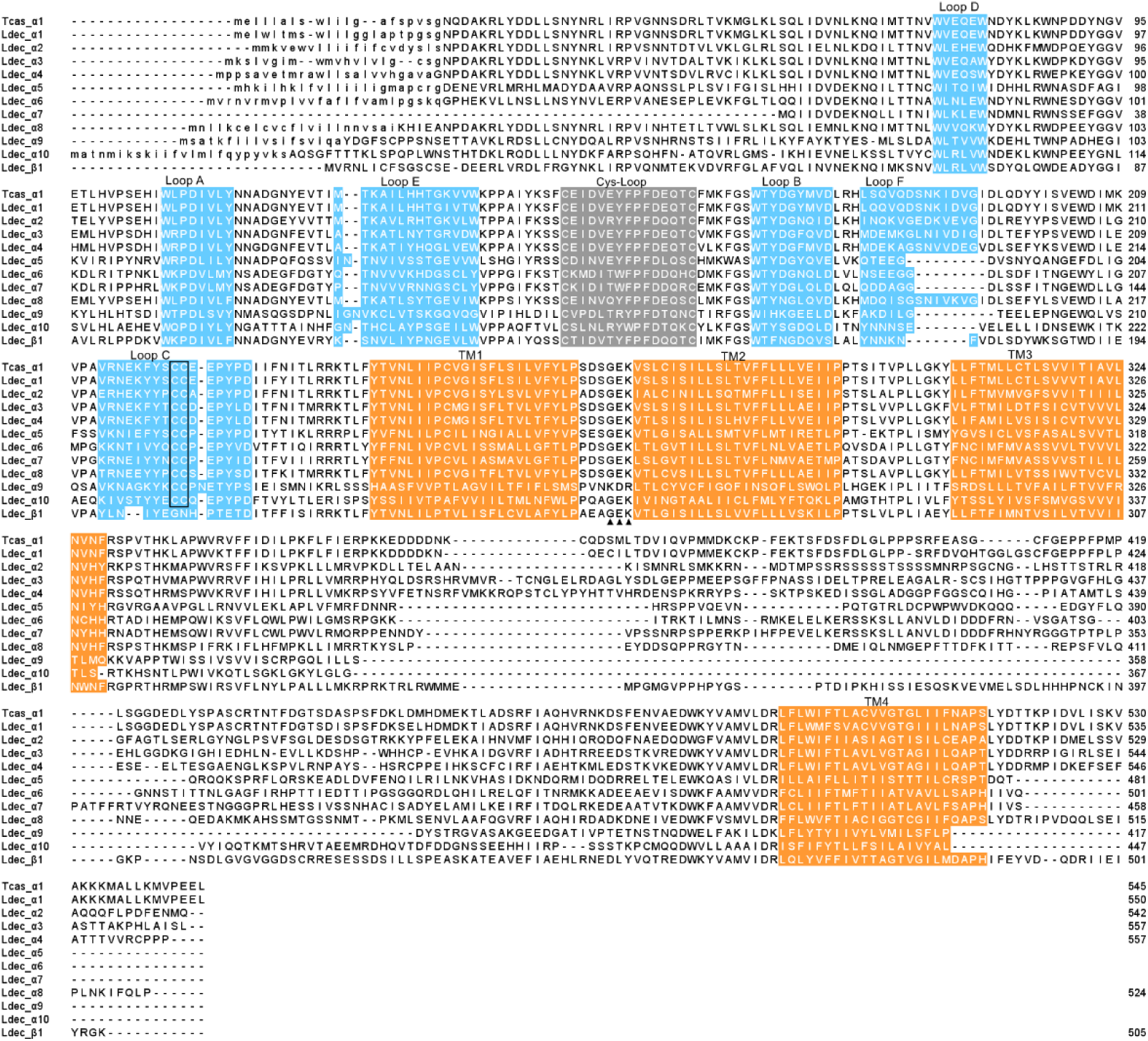
Protein sequence alignment of CPB nAChR subunits. The *T. castaneum* nAChR α1 was included as a reference (NCBI accession NP_001103245.1). N-terminal signal leader peptides are indicated by lowercase letters. Loops A-F are highlighted in blue; the cys-loops are highlighted in gray, and transmembrane domains are suggested by orange. The two consecutive cysteines in Loop C, characteristic of α subunits, are enclosed in a box. Three amino acids (GEK) preceding TM2 are indicated by solid triangles underneath.

**Table 3.** CPB nAChR subunits (identity/similarity) to *T. castaneum*.

|  | Ldec_a1 | Ldec_a2 | Ldec_a3 | Ldec_a4 | Ldec_a5 | Ldec_a6 | Ldec_a7 | Ldec_a8 | Ldec_a9 | Ldec_a10 | Ldec_b1 |
| --- | --- | --- | --- | --- | --- | --- | --- | --- | --- | --- | --- |
| Tcas_a1 | **90/94** | 53/69 | 55/66 | 52/65 | 29/48 | 33/51 | 34/52 | 53/69 | 15/32 | 18/37 | 38/56 |
| Tcas_a2 | 51/68 | **90/94** | 50/65 | 46/62 | 26/48 | 33/52 | 33/51 | 50/66 | 16/33 | 19/34 | 38/56 |
| Tcas_a3 | 55/67 | 50/65 | **87/92** | 67/76 | 28/48 | 32/49 | 34/51 | 55/67 | 15/32 | 19/34 | 41/59 |
| Tcas_a4 | 52/65 | 46/62 | 66/76 | **91/95** | 27/45 | 32/48 | 33/51 | 55/70 | 16/32 | 18/35 | 40/56 |
| Tcas_a5 | 23/40 | 21/40 | 23/40 | 22/38 | **63/67** | 26/41 | 25/41 | 21/38 | 13/27 | 15/28 | 24/40 |
| Tcas_a6 | 34/51 | 33/51 | 32/49 | 33/49 | 34/53 | **89/94** | 64/75 | 33/52 | 18/36 | 21/37 | 33/53 |
| Tcas_a7 | 33/51 | 34/52 | 33/49 | 34/50 | 32/51 | 62/74 | **89/94** | 33/51 | 16/33 | 19/37 | 32/51 |
| Tcas_a8 | 52/67 | 50/66 | 55/67 | 51/67 | 29/50 | 33/53 | 34/53 | **85/90** | 15/34 | 21/39 | 40/58 |
| Tcas_a9 | 15/30 | 15/32 | 15/32 | 14/31 | 16/36 | 15/34 | 14/33 | 14/32 | **43/66** | 23/47 | 14/33 |
| Tcas_a10 | 19/36 | 19/35 | 18/35 | 18/35 | 19/40 | 20/38 | 20/39 | 20/37 | 26/47 | **63/80** | 19/38 |
| Tcas_a11 | 52/66 | 50/66 | 55/66 | 52/66 | 28/48 | 33/51 | 33/50 | 84/90 | 15/34 | 21/38 | 39/57 |
| Tcas_b1 | 38/55 | 38/55 | 40/57 | 38/54 | 29/51 | 34/52 | 32/51 | 39/57 | 16/31 | 20/39 | **94/96** |

\*Potential orthologs are highlighted in bold.

In the nAChR amino acid sequence-based phylogeny (Figure 3), α1 - α8 and β1 subunits are relatively conserved across insect species [61]. The “nAChR α5 group” only includes non-dipteran orthologs, since dipteran α5 subunits were named following different nomenclatures [59]. For example, the *Bombyx mori* α5 shared 17% identity with *D. melanogaster* α5 [59]. Instead, *D. melanogaster* α5 subunits are clustered with non-dipteran α7 subunits, named here as “α7-like group”. In addition, two CPB nAChR, Ldec_α9, and Ldec_α10, clustered in the large divergent group.

**Figure 3.**
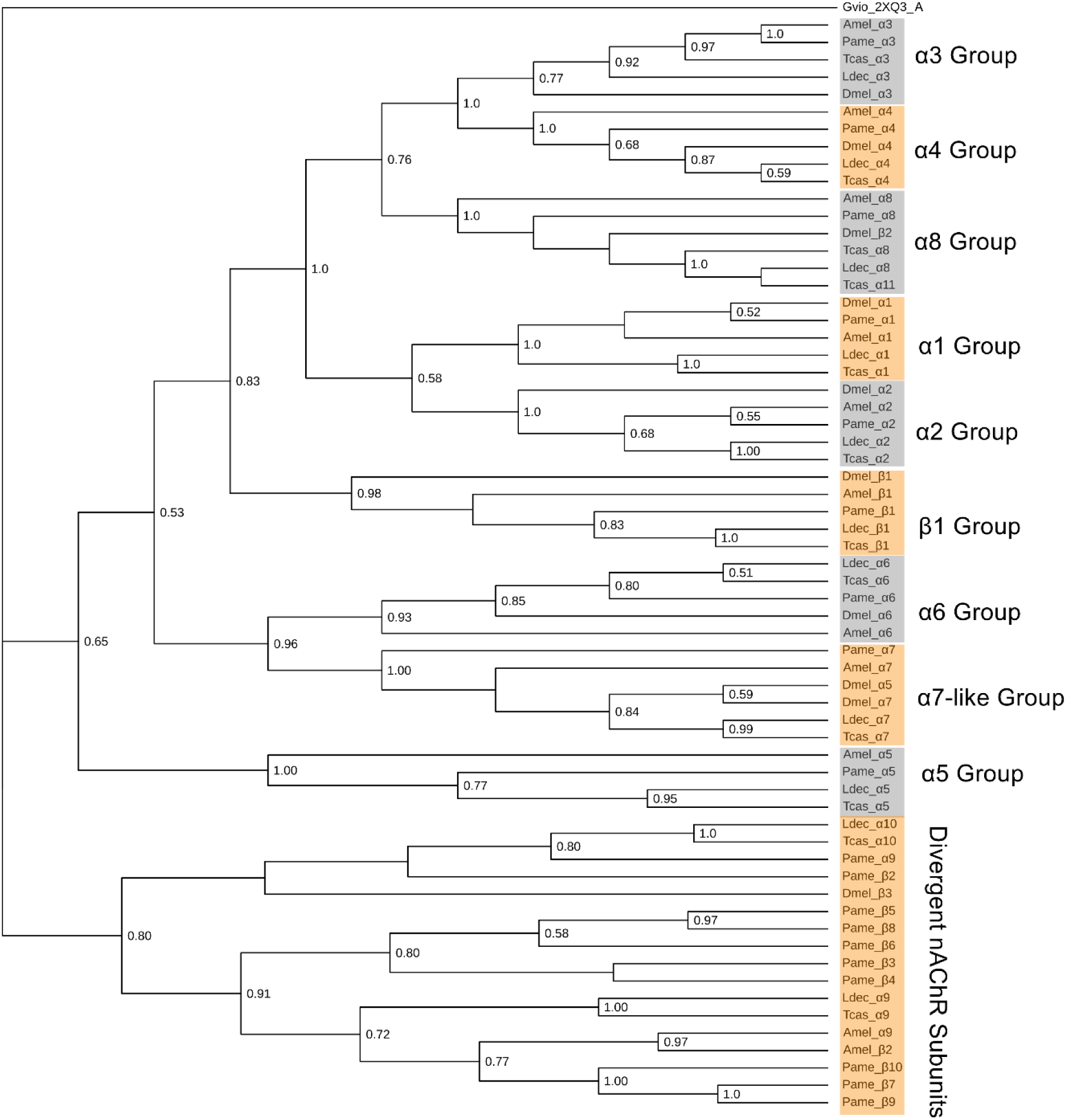
Phylogeny of nAChR subunits in five species. Species name annotations: Ldec-*L. decemlineata*; Dmel-*D. melanogaster*; Amel-*A. mellifera*; Tcas-*T. castaneum*; Pame-*Periplaneta americana*. Gvio_2XQ3_A, a bacterial ancestor of cysLGICs from *Gloebacter violaceus* was an outgroup, to which the tree was rooted. Bootstrapping values higher than 0.5 are displayed.

The *Ldec_*α*7* and *Ldec_β1* genes are located within 474 kb of each other on the X chromosome CM045700.1. Clustering of nAChR α7 and β1 subunits have similarly been observed in the genomes of *B. germanica, P. americana, A. mellifera, A. gambiae*, and *T. castaneum* [14, 19, 21, 62], suggesting conserved syntony and the possibility that their expression may be coordinated.

Preliminary evidence of alternative splicing in several subunits was supported by tBlastn results. Two alternatives were identified for *Ldec_α4* exon4, labelled exon4a and exon4b respectively (Figure 4a). Alternative splicing leads to amino acid polymorphism in agonist binding pocket forming region, Loop E and Loop B, which may contribute to the variable function of protein isoforms. The *Ldec_α6* has two potential alternatives in exon 8 (Figure 4b), encoding TM2, which forms the pore of the channel, possibly contributing to functional diversity [1]. Alternative splicing of exon 4 in the nAChR α4 and exon 8 in nAChR α6 has also been observed in other insect species [63].

**Figure 4.**
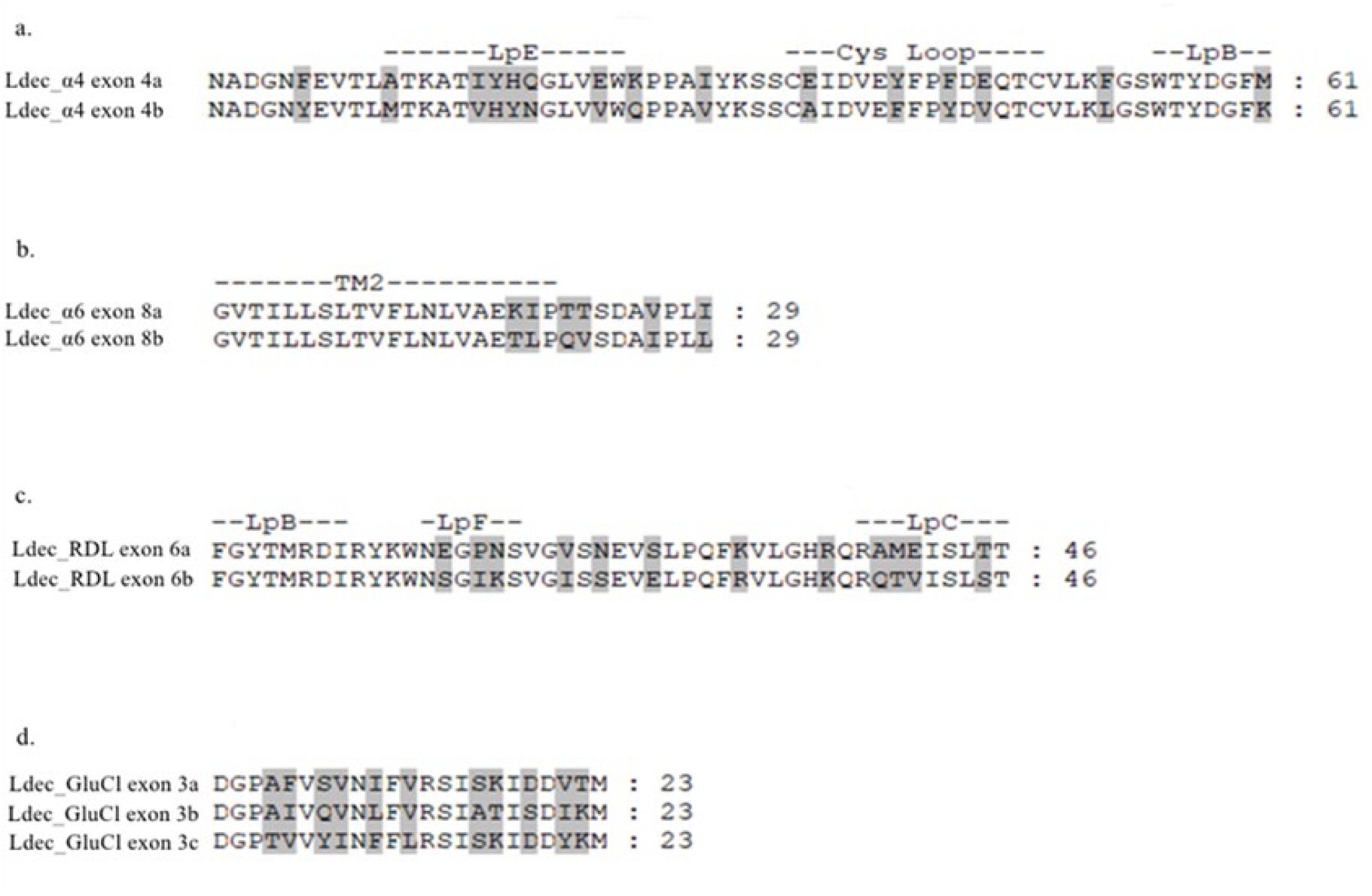
Potential alternative splicing. Variable sites are shaded. a) Ldec_a4 exon 4 splicing variants; b. Ldec_a6 exon eight splicing variants; c. Ldec_RDL exon six splicing variants; d. Ldec_GluCl exon three splicing variants.

A comparison of the CPB nAChR α6 gDNA and a transcript (GGNV01207620.1) sequence indicate three potential cases of adenosine (A) to guanosine (G) editing in gDNA sequence positions 27, 415, and 468 (Figure 5a). This conversion results in an S-to-L, N-to-D, and I-to-M amino acid substitution on protein positions 10, 139, and 156, respectively (Figure 5b). The S-to-L substitution may not affect the protein product since it is in the signal peptide region (refer to Figure 2), which will be trimmed in the final protein. The N-to-D substitution removed a potential N-glycosylation site, which may affect receptor maturation, channel desensitization, and conductance. It is worth noting that nucleotide differences between reference genomes may be due to genomic DNA variation rather than RNA A-to-I editing since these two sequences came from different CPB samples.

**Figure 5.**
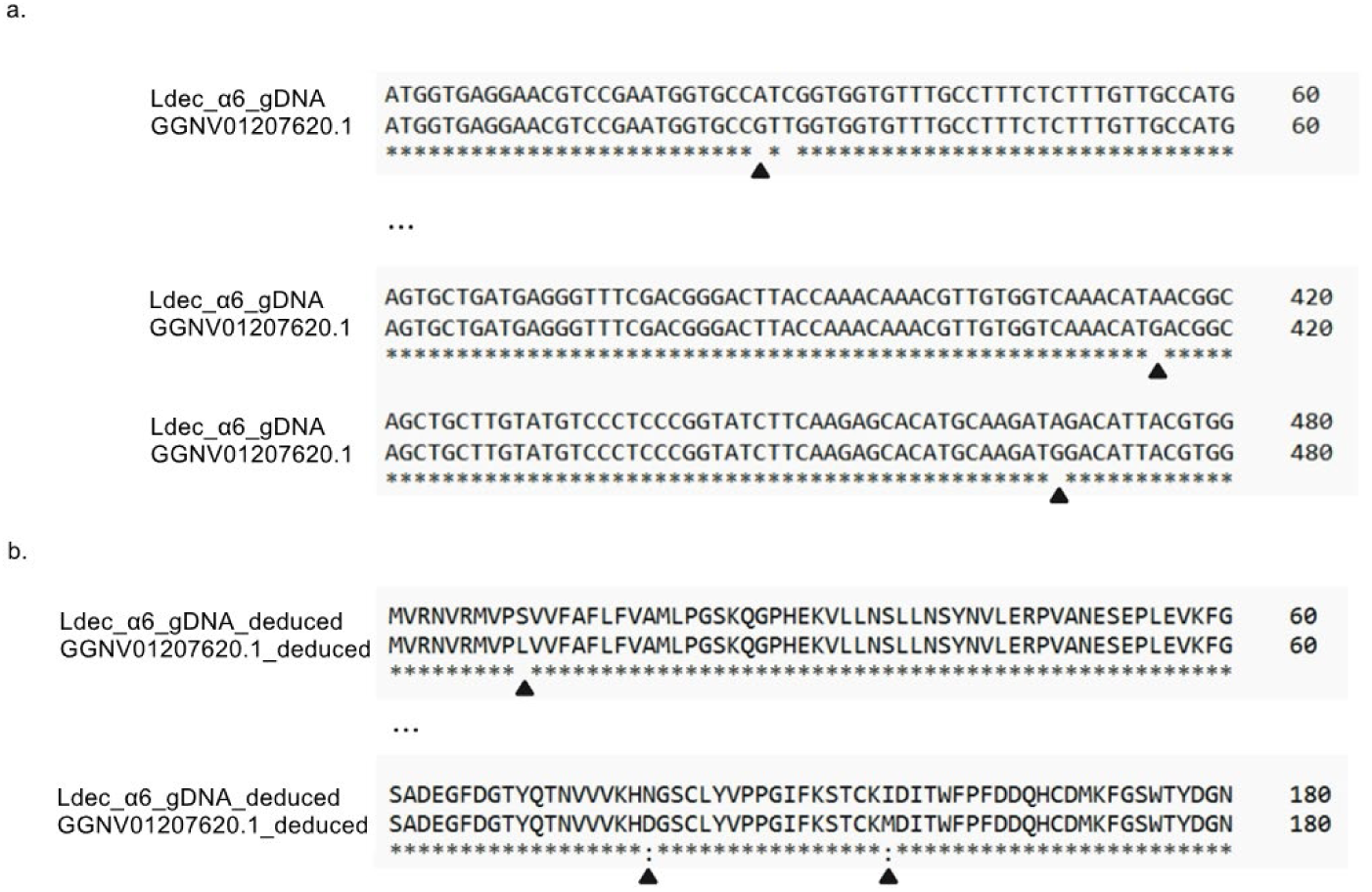
Potential RNA A-to-I editing on CPB nAChR α6. a). A-to-G conversion between the gDNA and the transcript at site 27, 415, and 468 respectively (indicated by solid triangles) b). Amino acid replacement due to RNA editing, indicated by solid triangles.

### GABA receptor subunits

In CPB, one ortholog of each GABA receptor subunit was identified and named Ldec_RDL, Ldec_GRD, and Ldec_LCCH3, respectively. The Ldec_RDL shows a high identity with its *T. castaneum* counterpart (Table 4) and an anion channel signature PAR motif preceding TM2 which is conserved across insect species (Figure 6). The Ldec_GRD and Ldec_LCCH3 lack the PAR motif (Figure 6), indicating their potential role as a cation channel. Two alternative exons 6, leading to 12 amino acid polymorphisms of the Ldec_RDL were detected by the tBlastn result (Figure 4c). Some of the amino acid variation occurs between loops F and C, within the ligand binding pocket, which may cause functional effects on receptors.

**Figure 6.**
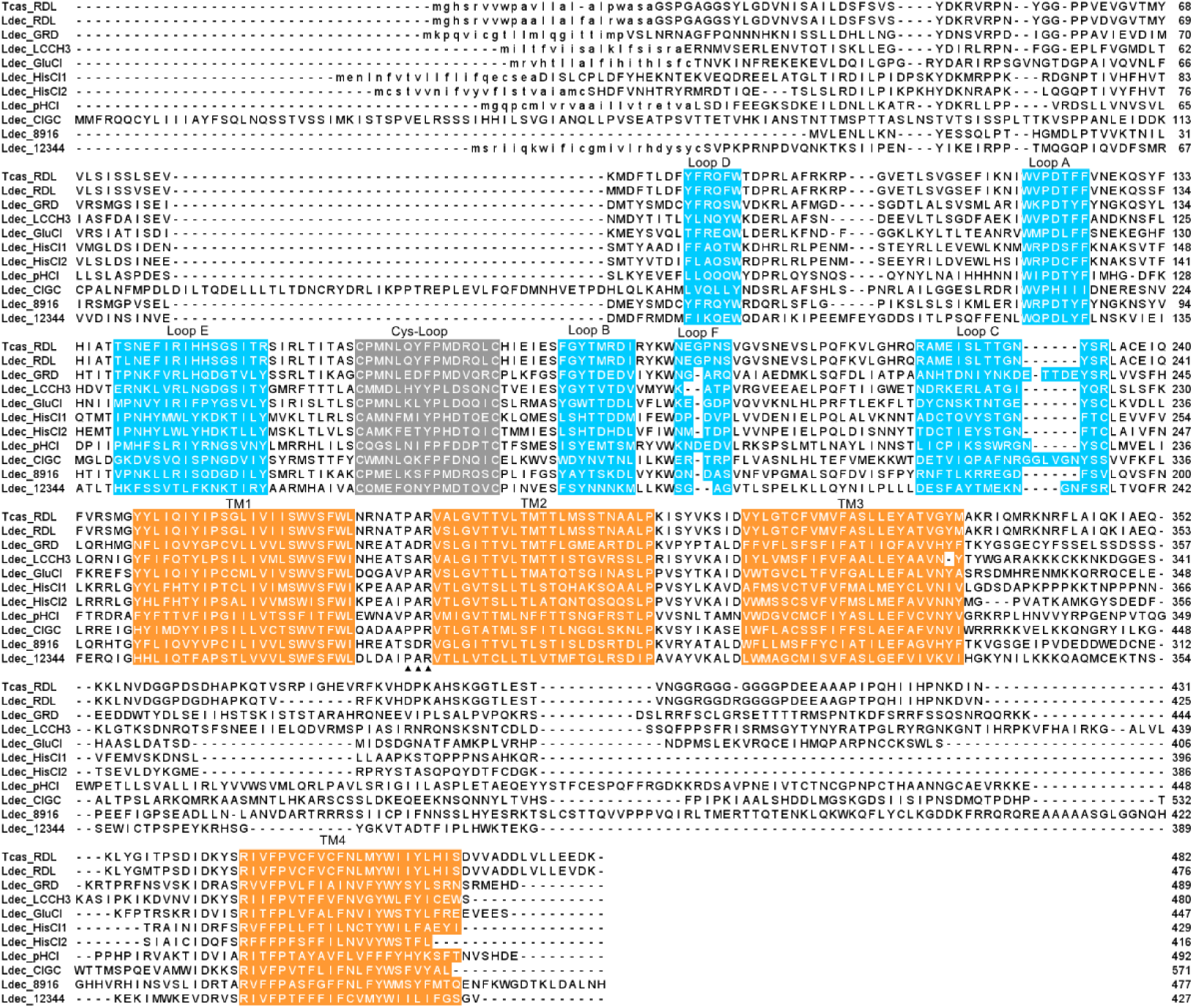
Protein sequence alignment of CPB cysLGIC subunits other than nAChR. *T. castaneum* RDL subunit (NCBI accession NP_001107809.1) is included as a reference. N-terminal signal leader peptides are indicated by lowercase letters. Loops A-F are highlighted in blue, the cys-loops are highlighted in gray, and transmembrane domains are suggested in orange. Three amino acids preceding TM2 are indicated by solid triangles underneath.

**Table 4.**
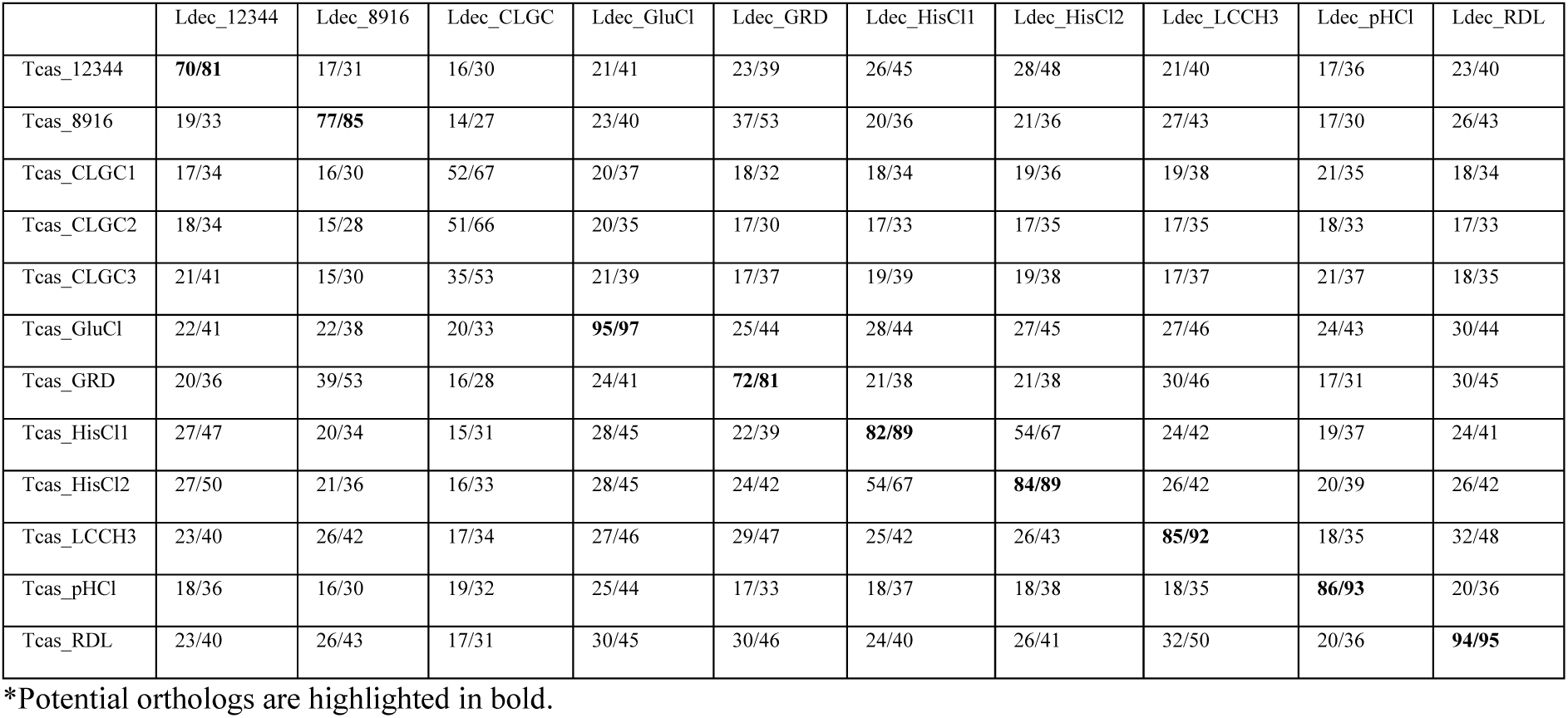
CPB non-nAChR cysLGIC subunits homology statistics (identity/similarity) to *Tribolium*.

|  | Ldec_12344 | Ldec_8916 | Ldec_CLGC | Ldec_GluCl | Ldec_GRD | Ldec_HisCl1 | Ldec_HisCl2 | Ldec_LCCH3 | Ldec_pHCl | Ldec_RDL |
| --- | --- | --- | --- | --- | --- | --- | --- | --- | --- | --- |
| Tcas_12344 | **70/81** | 17/31 | 16/30 | 21/41 | 23/39 | 26/45 | 28/48 | 21/40 | 17/36 | 23/40 |
| Tcas_8916 | 19/33 | **77/85** | 14/27 | 23/40 | 37/53 | 20/36 | 21/36 | 27/43 | 17/30 | 26/43 |
| Tcas_CLGC1 | 17/34 | 16/30 | 52/67 | 20/37 | 18/32 | 18/34 | 19/36 | 19/38 | 21/35 | 18/34 |
| Tcas_CLGC2 | 18/34 | 15/28 | 51/66 | 20/35 | 17/30 | 17/33 | 17/35 | 17/35 | 18/33 | 17/33 |
| Tcas_CLGC3 | 21/41 | 15/30 | 35/53 | 21/39 | 17/37 | 19/39 | 19/38 | 17/37 | 21/37 | 18/35 |
| Tcas_GluCl | 22/41 | 22/38 | 20/33 | **95/97** | 25/44 | 28/44 | 27/45 | 27/46 | 24/43 | 30/44 |
| Tcas_GRD | 20/36 | 39/53 | 16/28 | 24/41 | **72/81** | 21/38 | 21/38 | 30/46 | 17/31 | 30/45 |
| Tcas_HisCl1 | 27/47 | 20/34 | 15/31 | 28/45 | 22/39 | **82/89** | 54/67 | 24/42 | 19/37 | 24/41 |
| Tcas_HisCl2 | 27/50 | 21/36 | 16/33 | 28/45 | 24/42 | 54/67 | **84/89** | 26/42 | 20/39 | 26/42 |
| Tcas_LCCH3 | 23/40 | 26/42 | 17/34 | 27/46 | 29/47 | 25/42 | 26/43 | **85/92** | 18/35 | 32/48 |
| Tcas_pHCl | 18/36 | 16/30 | 19/32 | 25/44 | 17/33 | 18/37 | 18/38 | 18/35 | **86/93** | 20/36 |
| Tcas_RDL | 23/40 | 26/43 | 17/31 | 30/45 | 30/46 | 24/40 | 26/41 | 32/50 | 20/36 | **94/95** |

\*Potential orthologs are highlighted in bold.

### CPB glutamate and histamine-gated chloride ion channels

One glutamate ion channel gene (*Ldec_GluCl*) and two histamine-gated chloride ion channel genes (*Ldec_HisCl1*, and *Ldec_HisCl2*) were identified in the CPB genome. *T. castaneum* and CPB have high sequence similarity (over 82%) of these orthologs (Table 4). These three subunits display a PAR motif before TM2, consistent with their role as anion channels. Comparison among transcript-translated protein sequences indicates three alternative splicing products of exon 3 of the *Ldec_GluCl* gene (Figure 4d). Numbers of alternative GluCl exons 3 vary among species, with honeybees, fruit flies and two cockroach species having two [7, 19, 21] and the red flour beetles having three [14]. *Ldec_GluCl* exon 3 encodes the region proximate to loop D, which participates in the formation of ligand-binding pockets; therefore, amino acid polymorphisms due to alternative splicing may lead to variable ligand-binding properties among isoforms.

### CPB pH-sensitive chloride channel subunits

The Tcas_pHCl query matched two very similar, tandemly-arrayed CPB genomic regions on chromosome CM045712.1. Protein sequences deduced from these two genes are equal in length (426AA), having only one amino acid variant at position 356 (S/A), resulting in a 99.8% identity. This suggests that these two genes may be recently duplicated homologues of CPB pHCl. Therefore, they were named *Ldec_pHCl copy1* and *Ldec_pHCl_copy2* respectively (Table S2). However, only one CPB pHCl protein isoform was validated via transcript sequence and this isoform was used for phylogenetic analysis (Figure 7) and protein annotation (Figure 6).

**Figure 7.**
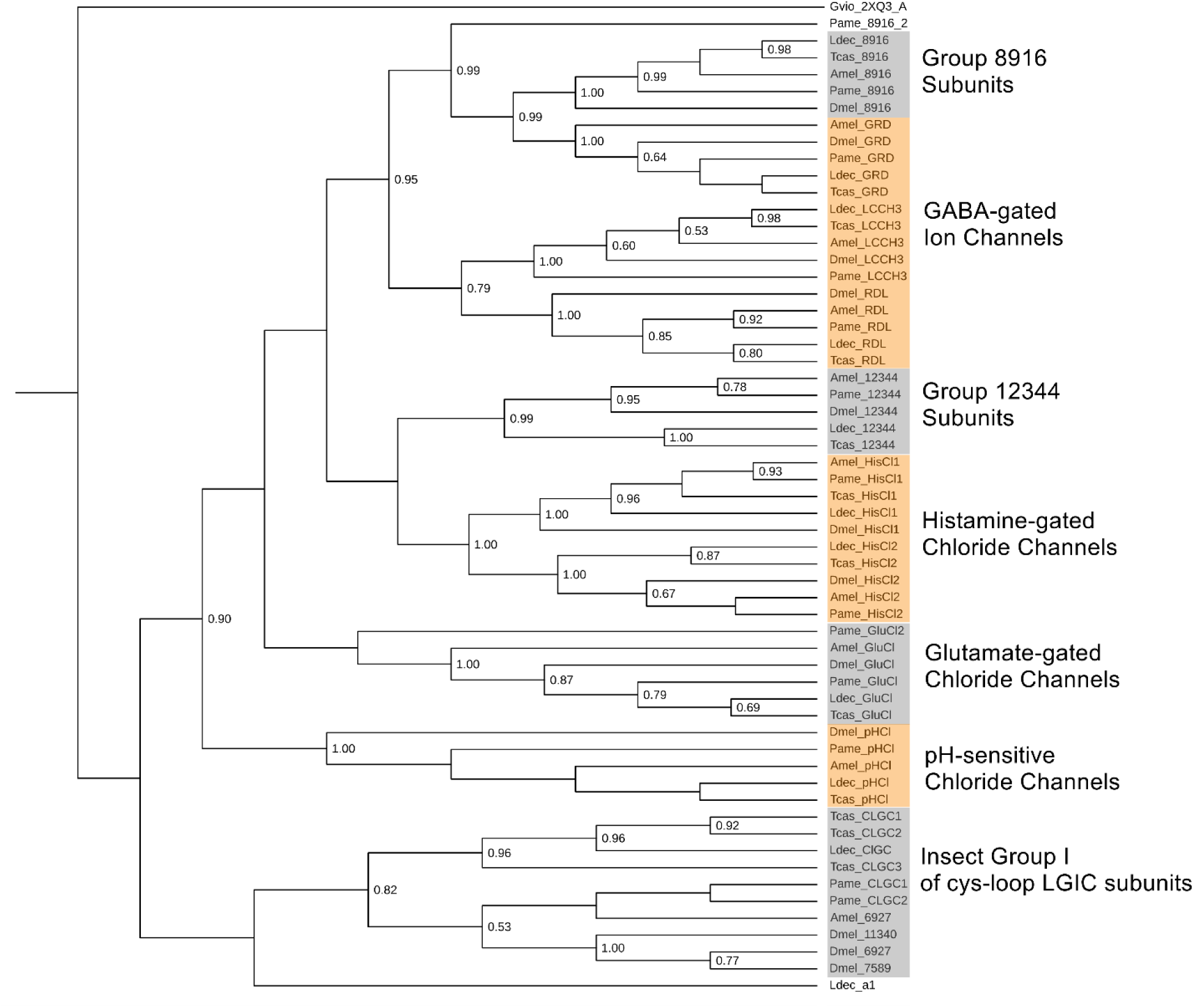
Phylogeny of non-nAChR cysLGIC subunits in five species. Gvio_2XQ3_A was an outgroup, and the tree was rooted in it. Ldec_α1 was also included as a reference. Bootstrapping values higher than 0.5 are displayed. Accession numbers of sequence used for phylogeny can refer to additional file 4: Table S4.

### Other CPB cysLGIC subunits

In CPB, only one gene belonging to Insect Group 1 cysLGICs was identified. Its position in the phylogeny reflects its relative similarity with three *T. castaneum* CIGCs (Figure 7). Therefore, it was named Ldec_CLGC. Insect Group 1 subunit homologs form an independent group in the phylogeny (Figure 7). [14][18]

Orthologs of Drosophila CG8916 and CG12344 were identified in the CPB genome and named Ldec_8916 and Ldec_12344, respectively. The Ldec_12344 displays a PAR motif before TM2 (Figure 6), indicating its potential role as an anion channel. The phylogeny shows that the Ldec_12344 and its orthologs grouped with HisCl subunits, while the 8916 group clustered with GABA-gated ion channel subunits (Figure 7).

## Discussion

We characterized the CPB cys-loop LGIC gene superfamily, which encodes for receptors that play major roles in the nervous system and are also targets of widely used insecticides. CPB has a similar number (22) of receptor subunit genes as other sequenced insects with greatest similarity to another beetle, *Tribolium*. Mechanisms increasing subunit diversity, such as alternative splicing and RNA A-to-I editing, are observed in nAChR α4, nAChR α6, RDL, and GluCl subunits.

### Ldec_α4, Ldec_α9 subunits of nAChR and Ldec_pHCl have unique features

Compared to other subunits, the Ldec_α9 subunit has low similarity with its *T. castaneum* counterpart, though *T. castaneum* remains the most similar of those included in the phylogram (Table 3). Ldec_α9 also lacks the GEK motif preceding TM2, which is usually conserved in nAChR subunits. To test whether the Ldec_α9 is properly categorized as a nAChR subunit we calculated the protein sequence identity between Ldec_α9 and all other CPB cysLGIC subunits. We observed that Ldec_α9 shares 9% to 14% identity with other CPB non-nAChR subunits, while its identity with other CPB nAChR can reach 25%. Further, the relative similarity to the *Tribolium* α9 in figure 3, indicates that Ldec_α9 is more likely a nAChR subunit than another non-nAChR cysLGIC subunit. Whether Ldec_α9 possesses the cation selectivity of other GEK-motif-containing nAChR subunits remains to be determined.

Of the nAChR subunits, α4 is the only one whose complete expression was not confirmed by either transcripts or cDNA. Figure 8 displays an alignment of all potential CPB nAChR α4 protein sequences deduced from different sources, including three transcripts obtained from a pest population in Wisconsin (GEEF01180990.1, GEEF01026894.1, and GEEF01038441.1), one cDNA from Xinjuang, China (KP090398.1) and the gDNA sequence (also from Xinjuang) used in this study, in addition to the *T. castaneum* nAChR α4 for reference. According to Tcas_α4 and the Ldec_α4 gDNA deduced sequence, a typical beetle nAChR α4 protein is expected to be over 550 amino acids; however, the longest transcript-deduced CPB protein (GGNV01187321.1_deduced) possesses only 182 amino acids. The cDNA-deduced sequence is slightly longer but still missing almost all N-terminal regions. This cDNA sequence was obtained in a previous study by mRNA RT-PCR, followed by a 5’- and 3’-Rapid amplification of cDNA ends (RACE) [50]. [59] Because of this, the short nAChR α4 cDNA may not be due to incomplete sequencing coverage or mis-amplification. These much shorter protein sequences deduced from transcripts and cDNA may indicate that CPB nAChR α4 are naturally truncated and since the cDNA and transcripts are from five different CPB populations, this truncated version of CPB nAChR α4 may be geographically widespread. Truncation of nAChR is also seen in other insect species, such as the truncated α4 and α5 transcripts found in *D*. *melanogaster* and *B*. *mori* [7, 59]. However, these truncated transcripts lead to a partial loss of transmembrane regions or ligand binding loops, rather than a nearly complete loss of N-terminal extracellular region seen in CPB nAChR α4. Whether this nAChR α4 truncation alters CPB’s natural biological processes and insecticide interactions remains to be determined as does the geographic and phylogenetic range of this truncation. Importantly, knocking out the nAChR α4 in a *D. melanogaster* strain led to a 1.491, 1.262, and 1.343-fold resistance ratios to sulfoxaflor, imidacloprid, and nitenpyram, respectively [64]. It is possible therefore that this structural change is in response to exposure of these diverse populations of CPB to neonicotinoid insecticides. Analysis of the roles of truncated nACh4 α4 subunits in the structure and function of nACh4 receptors would clearly be of value.

**Figure 8.**
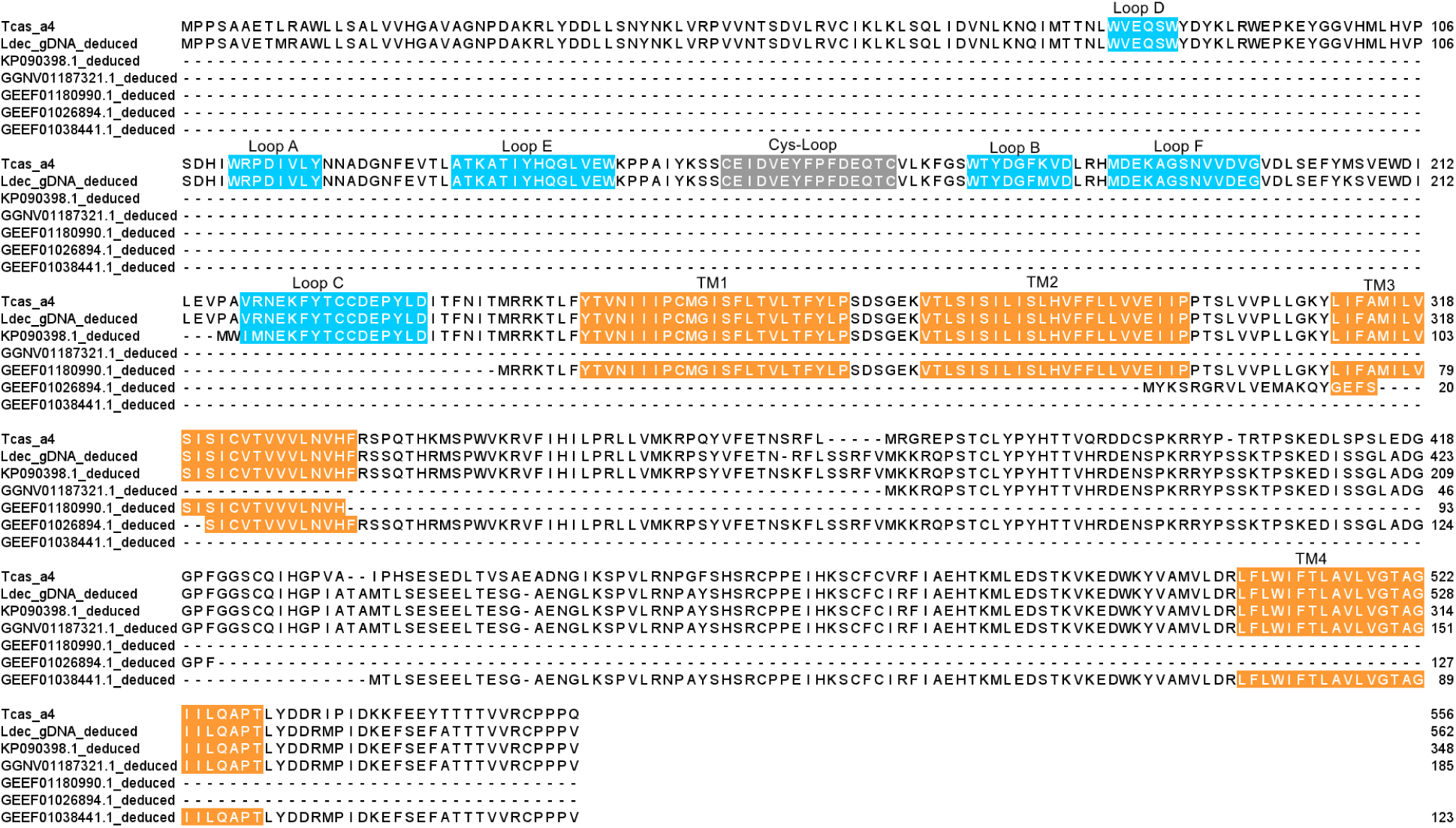
Protein sequence alignment of all potential CPB nAChR α4 isoforms. *T. castaneum* nAChR α4 (NCBI accession NP_001103246.1) is included for reference. Respectively, GGNV01187321.1, GEEF01180990.1, GEEF01026894.1, and GEEF01038441.1 are transcript sequences, KP090398.1 is a cDNA sequence.

For non-nAChR cysLGIC subunits, the expression of Ldec_RDL and Ldec_8916 were not validated by transcripts or database cDNA sequence. A previous study sought to amplify the CPB RDL mRNA using PCR [65] while only a fragmented product was obtained. The complete expression of these two subunits in the CPB population still needs to be validated.

We show preliminary evidence of the first *Ldec_pHCl* gene duplication observed in an insect species. This observation deserves additional validation because insect pHCl have potential as insecticide targets due to its exclusive occurrence in invertebrates [6].

### Chromosome level genome assembly indicates more genomic features

Improved CPB genome assemblies contributed here to revealing genomic features which, because genomic architecture has implications for adaptation to insecticide exposure, may provide insight into insecticide resistance [51]. To date, all known field-evolved neonicotinoid resistance-conferring mutations (not identified in CPB yet) are found on the β1 subunit, among which the R81T mutation is the most widely reported, detected in multiple *M. persicae* and *A. gossypii* populations worldwide [25–28, 66]. However, no homozygous 81TT individuals have yet been reported in any field population [25], possibly due to fitness costs associated with the 81TT genotype [67]. Colorado potato beetle is an XX-XO sex-determination system [68], of which the male individuals have a single X chromosome. Therefore, alleles present on the X chromosome of males would be functionally dominant for both resistance and fitness traits. If there were significant fitness costs of that allele, perhaps due to pleiotropic effects on neuronal function, this resistance mutation may not persist in CPB populations because of its deleterious effects on all male carriers of the resistance alleles.

Second, the improved completeness and contiguity of this newer genome assembly revealed the close physical proximity of some nAChR subunit genes. This may facilitate their coordinated expression and co-assembly into the native channel. For example, *D. melanogaster* nAChR α1, α2, and β2 subunit genes are clustered within 200kb of each other on chromosome 3; these three subunits were later shown to form robust functional nAChRs in *X. laevis* oocytes. Here, the protein-coding regions of *Ldec_α7* and *Ldec_β1* were observed to cluster within 470kb on the chromosome CM045700.1 (Table S2), suggesting that they are likely to co-assemble in native CPB nAChR.

## 5. Conclusion

CPB has a typical number of cysLGIC superfamily genes compared to other insect species and has members of all major groups. There is evidence that CPB has a recently duplicated pHCl gene, but further testing of this observation is needed. With the exception of one subunit with truncated transcripts (nAChR α4) CPB subunit genes were typical in structure as well. Evidence of alternative splicing and RNA editing were also identified in CPB subunit transcripts, potentially contributing to subunits’ structural and functional diversity. The characterization of CPB cysLGICs provides further insights into the evolution of this gene superfamily in insects. The sequence characterization provides an important basis for future studies of these ion channels as well as for the rational design of insecticides that control CPB, ideally with less effect on non-target organisms. Such studies can include the functional expression of certain subunits to determine their interaction features with native ligands and insecticides. In addition, based on the sequence characterization, RNA interference and gene editing techniques may also be applied to CPB nAChRs or other cysLGIC subunits to assess their role in neuronal function and response to insecticides.

## Supplementary information

Additional file 1: Table S1. A list of numbers of cysLGIC subtype genes in characterized insect species.

Additional file 2: Table S2. A list of potential CPB cysLGIC subunits and their chromosomal locations.

Additional file 3: Table S3. tBlastn results of gDNA extracted proteins against transcriptome and selection of protein sequence for alignment and phylogenic analysis.

Additional file 4: Table S4. List of NCBI GenBank accessions for other insect species used for phylogenetic analysis.

## List of abbreviations

ACh: Acetylcholinesterase
BLAST: Basic Local Alignment Search Tool
cDNA: complementary DNA
CLGC: Cys-loop Ligand Gated ion Channel
CNS: central nervous system
CPB: Colorado Potato Beetle
cysLGIC: Cys-loop ligand-gated ion channels
DNA: Deoxyribonucleic acid
GABA: Gamma-aminobutyric acid
gDNA: genomic DNA
GluCl: Glutamate-gated chloride channel
GRD: Gamma-aminobutyric acid receptor alpha-like
HisCl: Histamine-gated chloride receptors
IRAC: Insecticide Resistance Action Committee
LCCH3: Gamma-aminobutyric acid receptor subunit beta-like
mRNA: messenger ribonucleic acid
nAChR: nicotinic acetylcholine receptors
NCBI: National Center for Biotechnology Information
pHCl: pH-sensitive chloride channels
RACE: Rapid amplification of cDNA ends
RDL: Resistance to dieldrin
RNA: Ribonucleic acid
TM: Transmembrane

## Declarations

### Ethics approval and consent to participate

Not applicable

### Consent for publication

Not applicable

### Availability of data and materials

Links for the datasets supporting the conclusions of this article are included within the article and its additional files.

### Authors’ contributions

DC conceived the project, performed the research, and wrote the manuscript. DJH participated and critically revised the manuscript for important intellectual content. Both authors read and approved the final manuscript.

## Acknowledgments

Not finish yet

