## Supplemental Tables for "The cys-loop ligand-gated ion channel gene superfamily of the Colorado potato beetle, *Leptinotarsa decemlineata*"

Table S1. Numbers of cysLGIC superfamily gene in characterized insect species.

| | Dmel | Amel | Tcas | Nvit | Apis | Aaeg | Atum | Bger | Pame |
|---|---|---|---|---|---|---|---|---|---|
| **nAChR** | | | | | | | | | |
| α type | 7 | 9 | 11 | 12 | 9 | 10 | 11 | 10 | 9 |
| β type | 3 | 2 | 1 | 4 | 2 | 4 | 1 | 7 | 10 |
| **GABA** | | | | | | | | | |
| RDL | 1 | 1 | 1 | 1 | 2 | 1 | 1 | 1 | 1 |
| GRD | 1 | 1 | 1 | 1 | 0 | 1 | 1 | 1 | 1 |
| LCCH3 | 1 | 1 | 1 | 1 | 0 | 1 | 0 | 1 | 1 |
| **Others** | | | | | | | | | |
| 8916 | 1 | 1 | 1 | 1 | 0 | 0 | 1 | 2 | 2 |
| Insect group1 (CLGC) | 3 | 1 | 3 | 1 | 2 | 1 | 1 | 2 | 2 |
| HisCl | 2 | 2 | 2 | 2 | 2 | 2 | 2 | 2 | 2 |
| GluCl | 1 | 1 | 1 | 1 | 2 | 1 | 1 | 2 | 2 |
| pHCl | 1 | 1 | 1 | 1 | 2 | 1 | 1 | 1 | 1 |
| 12344 | 1 | 1 | 1 | 1 | 1 | 0 | 1 | 1 | 1 |
| **Total** | 22 | 21 | 24 | 26 | 22 | 22 | 21 | 30 | 32 |

Species name annotations: Dmel-D. melanogaster; Amel-A. mellifera; Tcas-T. castaneum; Nvit-Nasonia vitripennis; Apis-Acyrthosiphon pisum; Aaeg-Aedes aegypti; Atum-Aethina tumida; Bger-Blattella germanica; Pame-Periplaneta americana.

Table S2. Chromosomal location of CPB cysLGIC subunit genes

| Subunit gene | Chromosomal location | CDS start | CDS end |
|---|---|---|---|
| *Ldec_α1* | CM045695.1 | 67746761 | 67768291 |
| *Ldec_α2* | CM045696.1 | 3170836 | 3138756 |
| *Ldec_α3* | CM045703.1 | 13858834 | 13970425 |
| *Ldec_α4* | CM045695.1 | 41018307 | 41754641 |
| *Ldec_α5* | CM045698.1 | 63481056 | 63396375 |
| *Ldec_α6* | CM045700.1 | 61872075 | 62180805 |
| *Ldec_α7* | CM045700.1 | 49090340 | 49655565 |
| *Ldec_α8* | CM045702.1 | 41014845 | 40916109 |
| *Ldec_α9* | CM045705.1 | 44836427 | 44824104 |
| *Ldec_α10* | CM045702.1 | 7803128 | 7759587 |
| *Ldec_β1* | CM045700.1 | 49515645 | 49597496 |
| *Ldec_12344* | CM045702.1 | 23277189 | 23316372 |
| *Ldec_8916* | CM045698.1 | 41150513 | 41092358 |
| *Ldec_CLGC* | CM045706.1 | 4769722 | 4747013 |
| *Ldec_GluCl* | CM045700.1 | 16070642 | 16150978 |
| *Ldec_Grd* | CM045702.1 | 48892562 | 48874817 |
| *Ldec_HisCl1* | CM045702.1 | 53884993 | 53871120 |
| *Ldec_HisCl2* | CM045700.1 | 51999541 | 51966458 |
| *Ldec_Lcch3* | CM045698.1 | 41229977 | 41274666 |
| *Ldec_pHCl copy1* | CM045712.1 | 24506564 | 24420011 |
| *Ldec_pHCl copy2* | CM045712.1 | 25371690 | 25459705 |
| *Ldec_Rdl* | CM045709.1 | 22156746 | 22310306 |

Table S3. BLASTn results and the selection of protein sequence for alignment.

| Receptor Name | Subunit ID | Number of transcript hits with bit score > 900) | ID of the transcript producing the longest protein isoform | Completeness of "signature regions" on the longest isoform | Treatment for protein alignment |
|---|---|---|---|---|---|
| nicotinic Acetylcholine receptor | Ldec_α1 | 2 | GGNV01022290.1 | Complete | No treatment |
| | Ldec_α2 | 3 | GGNV01160150.1 | Complete | No treatment |
| | Ldec_α3 | 9 | GGNV01295170.1 | Complete | N/A |
| | Ldec_α4 | 3 | GGNV01187321.1 | Only TM4 present | Replaced with gDNA-translated protein |
| | Ldec_α5 | 1 | GGNV01170968.1 | Missing Loops D, A, E, and part of Loop B | Replaced with AWC68049.1 |
| | Ldec_α6 | 6 | GGNV01207620.1 | Complete | No treatment |
| | Ldec_α7 | 1 | GGNV01057173.1 | Complete | No treatment |
| | Ldec_α8 | 1 | GGNV01106564.1 | Complete | No treatment |
| | Ldec_α9 | 1 | GGNV01304972.1 | Complete | No treatment |
| | Ldec_α10 | 4 | GGNV01314871.1 | Complete | No treatment |
| | Ldec_β1 | 1 | GGNV01068309.1 | TM4 partially missing | Replaced by AKL79440.1 |
| GABA-gated ion channels | Ldec_RDL | NA | GGNV01394567.1 | Missing ligand binding loops and TMs | Replaced by XP_023012203.1 |
| | Ldec_GRD | 4 | GGNV01130196.1 | Complete | No treatment |
| | Ldec_LCCH3 | 2 | GGNV01135992.1 | Complete | No treatment |
| glutamate-gated chloride channels | Ldec_GluCl | 6 | GGNV01268739.1 | N-terminal loops missing | Replaced by XP_023018980.1 |
| Histamine-gated chloride channels | Ldec_HisCl1 | 3 | GGNV01338955.1 | Complete | No treatment |
| | Ldec_HisCl2 | 4 | GGNV01159996.1 | Complete | No treatment |
| pH-sensitive chloride channel | Ldec_pHCl_copy1 and copy2* | 2 | GGNV01044124.1 | Complete | No treatment |
| insect group 1 ligand-gated ion channels | Ldec_CLGC | 3 | GGNV01134008.1 | Complete | No treatment |
| Others | Ldec_8916 | 1 | GGNV01117992.1 | Missing Loops D, A, E, B, and F | Replaced by gDNA-translated protein |
| | Ldec_12344 | 1 | GGNV01048522.1 | Complete | No treatment |

XP_023012203.1 and XP_023018980.1 are predicted by automatic computational analysis; AKL79440.1 and AWC68049.1 are deduced from cDNA obtained by mRNA RT-PCR and 5'- & 3'- RACE.

*Protein sequence trasnslated from Ldec_pHCl_copy1 and Ldec_pHCl_copy2 only display one amino acid variance. GGNV01044124.1 were the best hitted transcript for both, and show a identical amino acid as Ldec_pHCL_copy2 at the polymophism site. Therefore protein sequence from GGNV01044124.1 was selected to represent both gene, and named Ldec_pHCl in the alignment.

Table S4 NCBI accession number of non-CPB cyLGIC sequences used for phylogeny.

| **Species** | **Subunits** | **NCBI Accesion Number** |
|---|---|---|
| *Apis mellifera* | Amel_α1 | NP_001091690.1 |
| | Amel_α2 | NP_001011625.1 |
| | Amel_α3 | NP_001073029.1 |
| | Amel_α4 | NP_001091691.1 |
| | Amel_α5 | AAS75781.1 |
| | Amel_α6 | AAY87894.1 |
| | Amel_α7 | NP_001011621.1 |
| | Amel_α8 | NP_001011575.1 |
| | Amel_α9 | NP_001091694.1 |
| | Amel_β1 | NP_001073028.1 |
| | Amel_β2 | NP_001091699.1 |
| | Amel_12344 | ABG75746.1 |
| | Amel_6927 | XP_006563765.1 |
| | Amel_8916 | NP_001071290.1 |
| | Amel_GRD | ABG75735.1 |
| | Amel_LCCH3 | NP_001071280.1 |
| | Amel_RDL | ABG75734.1 |
| | Amel_HisCl1 | NP_001071279.1 |
| | Amel_HisCl2 | ABG75740.1 |
| | Amel_GluCl | NP_001071277.1 |
| | Amel_pHCl | NP_001137350.1 |
| | Dmel_α1 | CAA30172.1 |
| | Dmel_α2 | NP_524482.1 |
| | Dmel_α3 | CAA75688.1 |
| | Dmel_α4 | CAB77445.1 |
| | Dmel_α5 | AAM13390.1 |
| | Dmel_α6 | NP_723494.2 |
| | Dmel_α7 | NP_001285436.1 |
| | Dmel_β1 | P04755.1 |
| | Dmel_β2 | CAA39211.1 |
| | Dmel_β3 | NP_525098.1 |
| | Dmel_12344 | NP_001286298.1 |
| | Dmel_6927 | NP_572194.1 |
| | Dmel_8916 | NP_573090.3 |
| | Dmel_11340 | NP_001303470.1 |
| | Dmel_7589 | NP_001261986.1 |
| | Dmel_GRD | NP_524131.1 |
| | Dmel_LCCH3 | NP_996469.1 |
| | Dmel_RDL | AAA28556.1 |
| | Dmel_HisCl1 | NP_524406.1 |

| | | |
|---|---|---|
| | Dmel_HisCl2 | NP_650116.2 |
| | Dmel_GluCl | AAG40735.1 |
| | Dmel_pHCl | NP_001034025.2 |
| | Pame_α1 | AFJ04793.1 |
| | Pame_α2 | AKV94621.1 |
| | Pame_α3 | AKR16132.1 |
| | Pame_α4 | AFA28129.1 |
| | Pame_α5 | GFCQ01005211.1 |
| | Pame_α6 | AEA40429.1 |
| | Pame_α7 | QQH14653.1 |
| | Pame_α8 | QQH14654.1 |
| | Pame_α9 | QQH14656.1 |
| | Pame_β1 | QQH14655.1 |
| | Pame_β2 | GFCQ01032711.1 |
| | Pame_β3 | GFCQ01027461.1 |
| | Pame_β4 | GAWS02039241.1 |
| | Pame_β5 | GFCQ01009686.1 |
| | Pame_β6 | GFCQ01010089.1 |
| | Pame_β7 | GFCQ01012153.1 |
| | Pame_β8 | GFCQ01034959.1 |
| | Pame_β9 | GBJC01015771.1 |
| | Pame_β10 | GFCQ01027794.1 |
| | Pame_12344 | GFCQ01007480.1 |
| | Pame_8916 | GFCQ01012789.1 |
| | Pame_8916_2 | MW206636 |
| | Pame_GRD | MW201215 |
| | Pame_LCCH3 | GFCQ01015543.1 |
| | Pame_RDL | BAW87781.1 |
| | Pame_HisCl1 | GFCQ01022534.1 |
| | Pame_HisCl2 | MW206637 |
| | Pame_GluCl | BAW87777 |
| | Pame_GluCl2 | GFCQ01031120.1 |
| | Pame_pHCl | GAWS02040818.1 |
| | Pame_GLGC1 | GAWS02050116.1 |
| | Pame_GLGC2 | GFCQ01015521.1 |
| | Tcas_α1 | NP_001103245.1 |
| | Tcas_α2 | NP_001103423.1 |
| | Tcas_α3 | NP_001107770.1 |
| | Tcas_α4 | NP_001103246.1 |
| | Tcas_α5 | NP_001103252.1 |
| | Tcas_α6 | NP_001107773.1 |
| | Tcas_α7 | NP_001103420.1 |

| | | |
|---|---|---|
| | Tcas_α8 | NP_001103419.1 |
| | Tcas_α9 | NP_001103424.1 |
| | Tcas_α10 | NP_001103247.1 |
| | Tcas_α11 | NP_001107771.1 |
| | Tcas_β1 | NP_001103418.1 |
| | Tcas_12344 | NP_001103243.1 |
| | Tcas_8916 | NP_001103425.1 |
| | Tcas_GRD | NP_001107772.1 |
| | Tcas_LCCH3 | NP_001103251.1 |
| | Tcas_RDL | NP_001107809.1 |
| | Tcas_HisCl1 | NP_001103422.1 |
| | Tcas_HisCl2 | NP_001103421.1 |
| | Tcas_GluCl | NP_001107775.1 |
| | Tcas_pHCl | NP_001107777.1 |
| | Tcas_GLGC1 | NP_001103250.1 |
| | Tcas_GLGC2 | NP_001103249.1 |
| | Tcas_GLGC3 | NP_001103248.1 |
